# CryoEM structures of the multimeric secreted NS1, a major factor for dengue hemorrhagic fever

**DOI:** 10.1101/2022.04.04.487075

**Authors:** Bo Shu, Justin S.G. Ooi, Aaron W.K Tan, Thiam-Seng Ng, Wanwisa Dejnirattisai, Juthathip Mongkolsapaya, Guntur Fibriansah, Jian Shi, Victor A. Kostyuchenko, Gavin Screaton, Shee-Mei Lok

## Abstract

Dengue virus infection can cause dengue hemorrhagic fever (DHF). Dengue NS1 is multifunctional: the intracellular dimeric NS1 (iNS1) forms part of the viral replication complex, the extracellular multi-oligomeric secreted NS1 (sNS1) is a major factor contributing to DHF. The structure of the iNS1 is well studied but not sNS1. Here we show the tetrameric (stable and loose conformation) and hexameric structures of sNS1. Stability of the stable and loose tetramers is determined by the conformation of their N-terminal domain – elongated β- sheet or β-roll. Binding of an anti-NS1 Fab breaks the loose tetrameric and hexameric sNS1 into dimers, whereas the stable tetramer remains largely unbound. Our results show detailed quaternary organization of different oligomeric states of sNS1 and will contribute towards the design of dengue therapeutics.

## Main

Dengue is a positive sense single-stranded RNA enveloped virus. It belongs to the family *Flaviviridae*, which also includes significant human pathogens like West Nile (WNV), yellow fever (YF) and Zika virus (ZIKV). Dengue virus causes disease that ranges from mild dengue fever to the severe dengue hemorrhagic fever (DHF) and the dengue shock syndrome (DSS)^1^. There are four DENV serotypes. Dengvaxia, which is currently the only licensed dengue vaccine, has an overall efficacy of only 45%, and there is a possibility of priming the previously immune naïve children to develop severe disease in subsequent natural infection after vaccination ^2^. It has proven to be difficult to develop an effective vaccine using whole dengue virus or its surface E protein, as the vaccine would need to stimulate equally strong immune responses simultaneously against all four dengue serotypes or else it can lead to a possibility of antibody-dependent enhancement (ADE) of infection^3^. This could lead to severe DHF/DSS.

Recently, dengue non-structural protein 1 (NS1) has been investigated as an alternative protein candidate for vaccine development. It has been shown to play an important role in causing the vascular leakage symptoms in DHF/DSS^4^. NS1 is secreted alongside dengue virus particles during an infection. It is a multifunctional protein which exists in different oligomeric forms intracellularly and extracellularly^5^, and they play distinctly different roles. The intracellular NS1 (iNS1), exists as membrane associated dimeric form and is a part of the viral replication complex. The extracellular secreted NS1 (sNS1), exists in higher oligomeric forms. sNS1 likely causes vascular leakage symptoms in DHF/DSS via two pathways – indirectly, by stimulating the innate immune response through activation of complement and Toll-like receptors^6^, and directly, by binding to endothelial cells causing endothelial hyperpermeability. sNS1 when bound directly to endothelial cells causes barrier dysfunction of the cell monolayer as demonstrated in an *in vitro* trans-endothelial electrical resistance (TEER) assay^4,7^. *In vivo* studies show that the sNS1 is present at very high concentrations in patient sera and correlates closely with the severity of disease ^8^. Mice injected with sNS1 along with a sublethal dose of DENV were observed to succumb to infection. Injection of sNS1 alone also causes vascular leakage in mice^4^. NS1-immune polyclonal serum and anti-NS1 monoclonal antibodies (MAbs) can protect mice against lethal DENV2 challenge ^4^. sNS1 was also found to enhance viral infection in mosquitoes by downregulating mosquito midgut immune genes^9^. sNS1 is therefore an important viral protein both for viral infection (mammalian cells and mosquitoes) and pathogenesis, and it is important to understand the structure of the sNS1 protein.

iNS1 (45 kDa) is first made in the endoplasmic reticulum (ER) as a monomer. It then forms dimers and is glycosylated in the ER and the trans-Golgi network. It is eventually transported to the cell membrane where some is secreted outside the cell, becoming sNS1. There is a high-resolution crystal structure of full length dimeric dengue iNS1^10^ produced by baculovirus expression and subsequent extraction from cellular membrane compartments. There are also crystal structures of the dimeric form of NS1 protein from other flaviviruses^11^ and they are largely similar to that of the DENV iNS1. Each protomer of the iNS1 contains three domains (Extended data Fig. 1A): 1) the N-terminus contains the β-roll domain (amino acids 1-30) that interacts with the same domain from the other protomer within the dimer to form a roll-like structure (named β-roll). This domain is generally hydrophobic and is thought to interact with cellular membranes. 2) Following the β-roll domain is the wing domain (amino acids 31-181). The wing domain is the most flexible domain as it differs between different flavivirus NS1 crystal structures ^11^. 3) the β-ladder domain (amino acids 182-352), consists of a continuous arrangement of 10 β-strands. The β-ladder domains from the two iNS1 protomers line up with each other to form a long ladder-like pattern. One side of the β-ladder faces the hydrophobic β-roll, while the other side is decorated with its highly charged spaghetti loops. The interaction between protomers within the dimer is stabilized by a large number of hydrogen bonds^12^.

About a decade ago, two very low resolution (23 Å and 30 Å) EM structures of sNS1 were determined: a cryoEM structure reconstructed from images acquired by using an 120kV LaB_6_-equipped electron microscope (Philip CM12 TEM)^13^ and the other, a negative stained room temperature EM structure (Extended data Fig. 1B)^14^. Both maps had an overall hollow barrel-like structure lacking fine structural details. Since then, the availability of 300kV transmission electron microscopes equipped with direct electron detectors and field emission gun electron sources, as well as improved cryoEM reconstruction methods, have made more routine determination of cryoEM structures of small proteins to high resolution possible. Here we report the cryoEM structures of DENV2 sNS1 tetramers (dimer of dimers) and hexamers (trimer of dimers). We observe sNS1 exists predominantly in the tetrameric form. There are two tetrameric states - stable and loose, and some of these structures are determined to ∼3.5Å resolution. The most obvious and dramatic difference between the stable and loose tetramers is the organization of their N-terminal domains – elongated β-sheet and β-roll, respectively. Only a minority of the sNS1 population is hexameric and this cryoEM structure was determined to 8Å resolution. We also determined a structure of sNS1 complexed with a Fab 5E3 to 3.5Å resolution. Most of the sNS1 (loose tetramers and hexamers) dissociates into dimers when bound to Fab, while the stable tetramer remains largely unbound. This is consistent with our results showing the ability of the Fab 5E3 to only partially inhibit sNS1-induced endothelial cell permeability. This study shows sNS1 exists in a heterogenous mixture of oligomerization states, and we present the first high resolution structures of sNS1.

### sNS1 from DENV2 contains heterogenous oligomeric states with tetramers as the predominant population

His-tagged NS1 was expressed in human embryonic kidney (HEK) cells. We harvested the sNS1 from tissue culture supernatant and after purification, we analyzed it using coomassie blue stained semi-denaturing SDS-PAGE gel (the sample was not boiled nor exposed to DTT). The gel shows clear bands corresponding to dimers and tetramers (Fig. 1A). As the sample was exposed to SDS, a detergent, the presence of tetramer band suggests that the tetramer is resistant to detergent. We then performed cross-linking of sNS1 with bis(sulfosuccinimidyl) suberate (BS^3^) before analyzing it by denaturing SDS-PAGE. This showed the presence of aggregates, which could be due to BS^3^ non-specifically crosslinking proteins, as well as bands for hexamer, tetramer, dimer and monomer (Fig. 1B). These results also showed that the tetrameric form is the predominant oligomerization state in the NS1 protein sample.

**Fig. 1.**
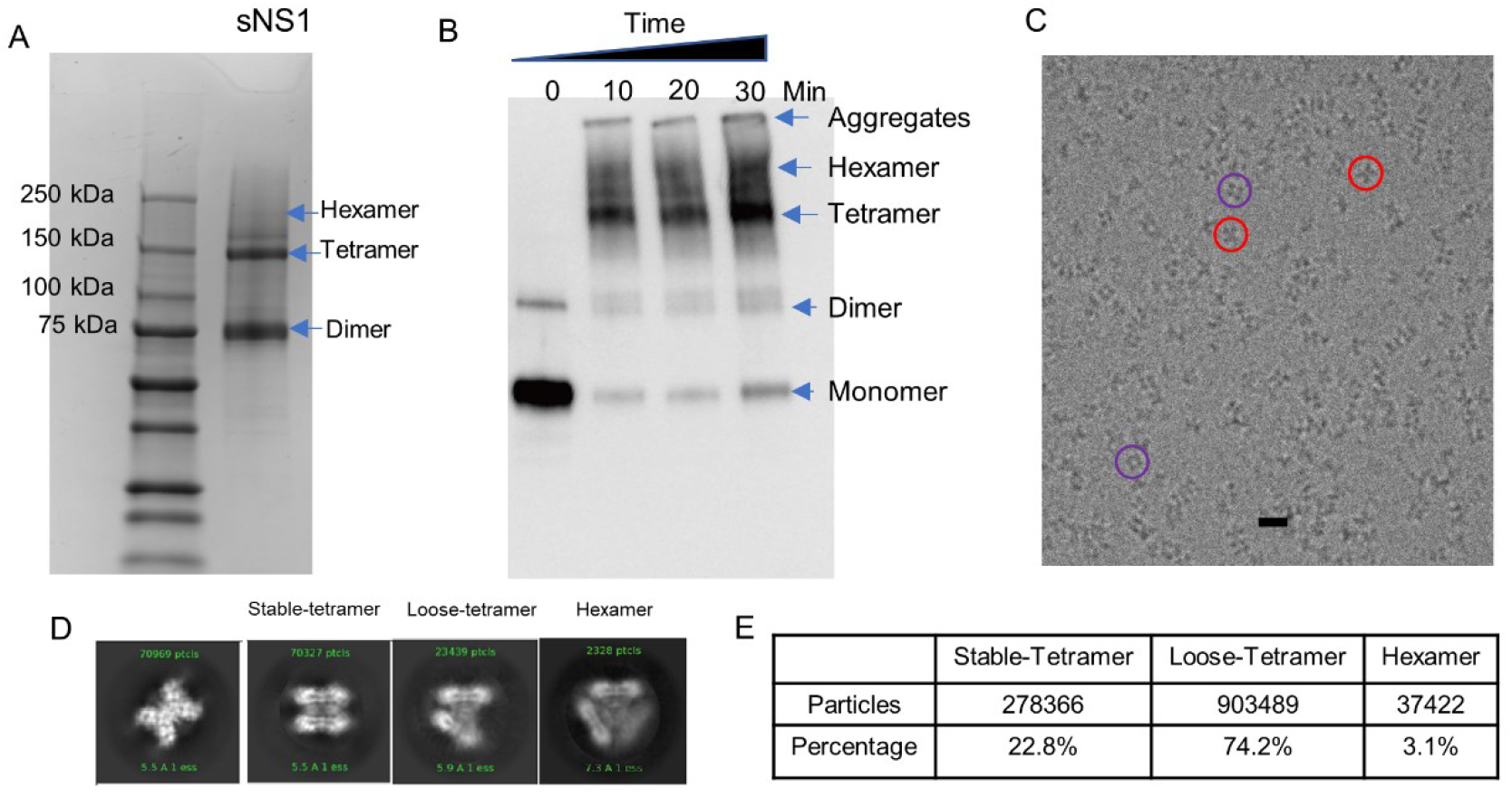
Profile of the DENV2 sNS1. **(A)** Semi-denaturing SDS-PAGE gel (non-boiling and no DTT) shows the clear dimer and tetramer bands, along with a very faint hexamer band. **(B)** Western blot denaturing SDS-PAGE gel (using Anti-polyHistidine−Peroxidase antibody) analysis of sNS1 cross-linked with BS^3^ at different incubation times. Aggregates, hexamers, tetramers, dimers and monomers were seen. (**C)** A cryoEM micrograph of the sNS1 showing distinct-shaped particles, e.g., butterfly-shaped (red circles) and cube-shaped particles (purple circles). Scale bar is 10 nm. (**D)** The 2D class averages indicate that the particles are heterogenous, with some likely existing as tetramers (stable and loose forms) and others as hexamers. (**E)** The distribution of particles in different oligomerization states determined after 3D classification and refinement. Most of the particles are tetramers (loose and stable) and hexamers are a minority in the population.

We collected 7714 cryoEM micrographs and observed particles with distinct shapes (Fig. 1C). We then performed several cycles of 2D classification and alignment of the boxed particles. The 2D class averages show particles with different features (Fig. 1D) - some appearing like stable and loose tetramers and others like hexamers. We did not observe any sNS1 dimers. This suggests that sNS1 consists of heterogenous populations of particles with likely tetrameric and hexameric states.

### CryoEM reconstruction of different oligomeric states of sNS1

As sNS1 consists of a heterogenous population of particles of different oligomerization states, we performed a 3D classification using the program Relion ^15^. The first round of unsupervised classification was done assuming no symmetry (C1) to avoid bias towards any specific oligomerization states. We then performed rough fitting with a dimeric NS1 crystal structure into these maps to determine their oligomerization states (Extended data Fig. 2). We observed two maps that correspond to tetramers: one with clear densities suggesting that the overall structure is quite stable (Extended data Figs. 2), while the other with looser interactions between the two dimers and poorer map resolution. We refer the former as stable tetramer and the latter, as loose tetramer. We also observed a 3D class that is hexameric in structure. Consistent with the 2D class averages, the 3D classification also did not yield any dimeric structures. We then conducted further classification and refinement. For the stable tetramers, we imposed D2 symmetry and obtained a final map of 3.5Å resolution (Fig. 2A), as measured by gold standard FSC curve cutoff at 0.143. For the loose tetramers, no symmetry was imposed, and we determined one subclass of the loose tetramer to 8 Å resolution (Fig. 3A(i) and Extended data Fig. 4A). We then performed local refinement on one of its dimers and obtained a 3.4 Å cryoEM density map (Fig. 3A(ii) and Extended data Fig. 4B). For the hexamers, since we only had a limited number of particles (∼30,000 particles), we obtained an 8Å map (Figs. 3B and Extended data Fig. 5).

**Fig. 2.**
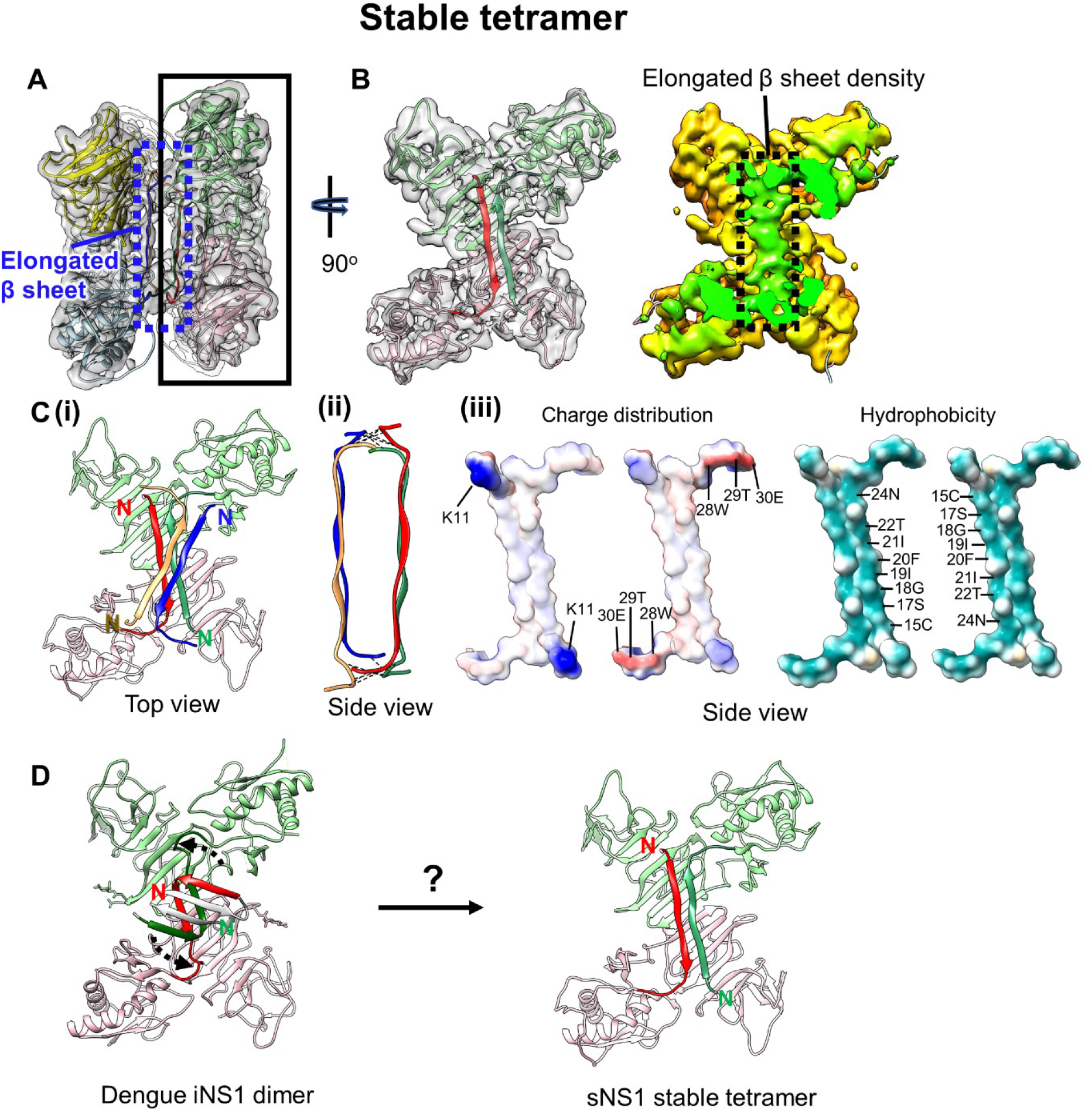
Structure of the stable sNS1 tetramer. (**A)** Two NS1 dimeric structures fitted into the cryoEM map (reduced sharpening was applied to make the density of the elongated β-sheet clearer). One dimer is colored in pink and light green and the other in yellow and light blue. **(B)** Left: the overall structure of the sNS1 dimer is similar to that of iNS1 except for the β-roll region (residues 1-30) in iNS1, which is an elongated β-sheet (residues 15-24) in sNS1. View from the elongated β-sheet of the black boxed region in (A). The respective elongated β-strand of each protomer is highlighted in a darker shade of the same color (red and dark green). Right: surface density map with the regions corresponding to the elongated β-sheet colored in green, and the rest of the map in yellow. **(C)** Interaction of the elongated β-strands between the two dimers (red/green and yellow/blue). (i) Top view showing the elongated β-sheets of the red/green dimer interacting with that of the yellow/blue dimer (only its elongated β-sheet is shown). (ii) Side view showing possible hydrogen bonds or electrostatic interactions between the two elongated β-sheets from opposite dimers (dotted black lines). These interactions are identified by the distance between Cα backbone (< 8Å) and also their charge characteristics. (iii) The interacting residues are complementary in both charges and hydrophobicity. Open book representation of (ii): (Left) charge, and (Right) hydrophobicity distributions on the surface of the two elongated β-sheets. For charge, positive charges are colored in blue while negative charges are red. For hydrophobicity, cyan color indicates hydrophobic residues. Possible interacting residues are labelled. **(D)** Comparison of iNS1 β-roll with sNS1 stable tetramer elongated β sheet and how it might change from the β-roll structure (dotted arrow on the iNS1 structure) into the elongated β sheet structure.

**Fig. 3.**
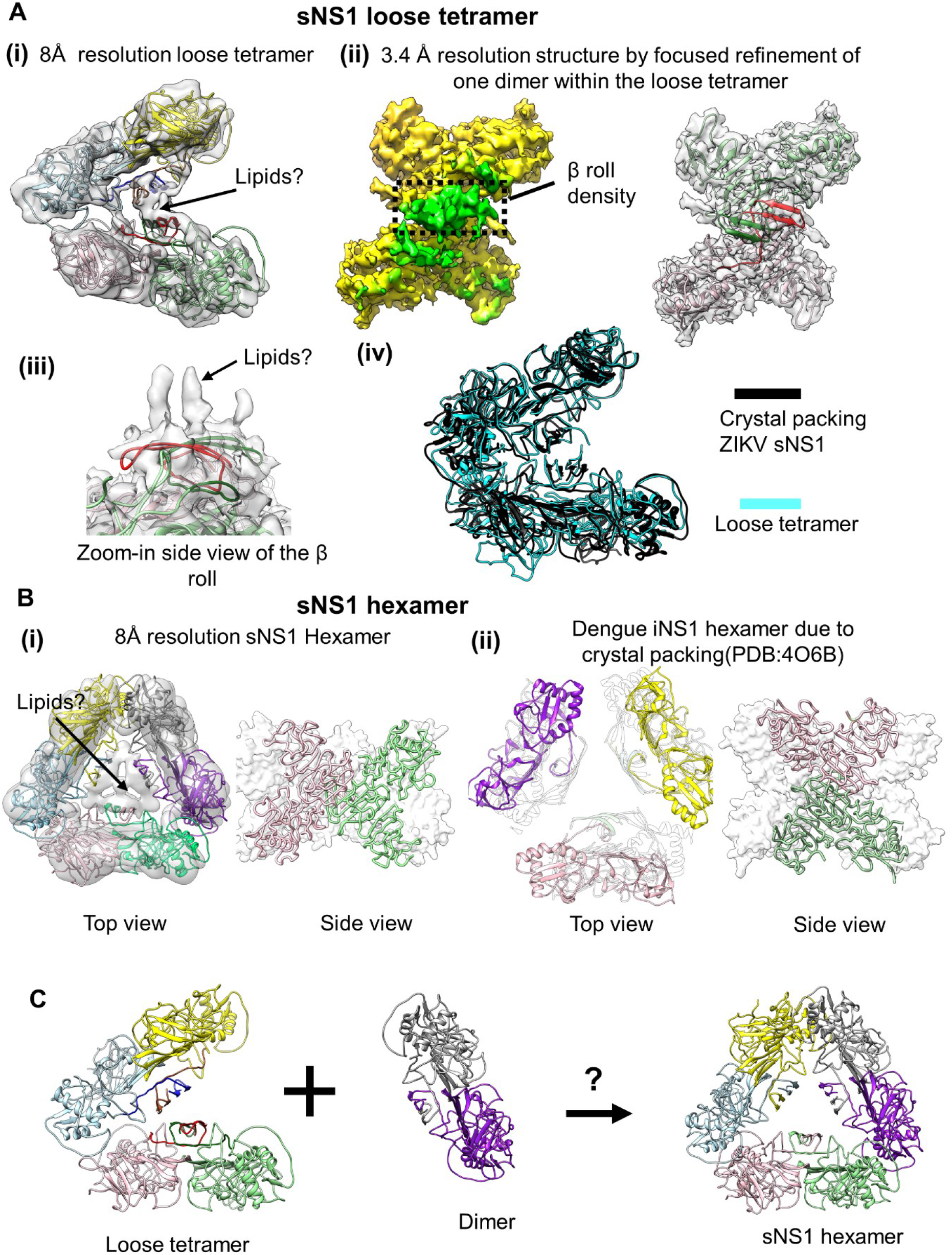
CryoEM structures of the sNS1 loose tetramer and hexamer. **(A)** (i) The fit of sNS1 into the 8 Å resolution loose tetramer density map. (ii) Focused refinement of one dimer of the loose tetramer yields a 3.4Å resolution map. An iNS1 dimer fits well into the density showing that the loose tetramer contains a β-roll structure similar to that of iNS1. Left: cryoEM map with the density corresponding to the β-roll colored in green and the rest in yellow. Right: the fit of the red/green dimer into the density. (iii) Zoom-in side view of the β-roll density (from (ii)) showing uninterpreted density that might belong to lipid molecules. (iv) Superposition of our loose tetramer with the tetramer observed in the crystal packing of ZIKV sNS1 structure (PDB: 5GS6) shows that they have similar structures. **(B)** Comparison of our sNS1 hexamer with the dengue iNS1 hexamer from the crystal packing. (i) Top and side views of the 8Å resolution sNS1 hexamer structure. Left: fit of three dimers into the cryoEM map. We are unable to discern whether sNS1 adopts a β-roll or an elongated β-sheet structure, as the interior of the hexamer contains unfeatured density. It may also contain lipid cargo as suggested by Gutsche and colleagues ^14^ (ii) The previously published dengue iNS1 structure ^10^ shows hexamers due to crystal packing; however, there is little interaction between the iNS1 dimers (left; top view). The dimers in the hexamer (right; side view) are also oriented differently to our hexameric structure (in (i), side view). **(C)** Interaction of a loose tetramer with a dimer could lead to the formation of a hexamer.

### Structure of sNS1 stable tetramers

The 3.5Å resolution sNS1 stable tetramer cryoEM map shows clear separation between β-strands (Extended data Figs. 3C-D). We also calculated the local resolution (Extended data Fig. 3A) of the map. While most of the density map is ∼3.5 Å resolution, parts of the map (corresponding to the β-ladder) are at ∼3.0Å resolution with some of the side chain densities resolved (Extended data Fig. 3D). The resolution of the wing domain and the β-roll was lower (4.0 to 4.5 Å). The density of both N- (residues 1-11) and C- terminal ends (residues 341-352) is missing from the map.

We interpreted the stable tetramer (dimer of dimers) cryo-EM density by fitting in the zika dimeric NS1 crystal structure (PDB:5GS6), because the structure is more complete than the dengue iNS1, and then mutated it into dengue NS1 sequence (Figs. 2A-B and Extended data Fig.3B). After fitting, we compared one of our sNS1 dimer in the stable tetramer to the dengue iNS1 dimer (PDB:4O6B) and it shows the wing domain of our sNS1 dimer is rotated by 6.8° with respect to the β-ladder. The β-roll structures of the two dengue iNS1 dimers cannot fit well into the cryoEM density of the stable tetramer and they clash with each other. Indeed, the space (25Å) between two dimers within the stable tetramers is not big enough to accommodate the two iNS1 β-rolls. We observed that the β-roll densities, instead of concentrating at the middle of the dimers as observed in iNS1 (Extended data Fig. 1), have an elongated shape that stretches across the two protomers within a dimer (Figs. 2A and B). This suggests that they have very different tertiary structures. Fitting of the density map shows that residues 15 to 24 form a long β-strand which pairs with the same β-strand of the other protomer within the dimer to form a long anti-parallel β-sheet (hereafter, referred to as elongated β-sheet) (Figs 2A-B). As the resolution of this region is ∼4.5 Å, we analyzed their interaction by using a distance cut off of 8 Å between the Cα backbone. Using this criterion, we observed that this elongated β-sheet from one dimer likely interacts with the same β-sheet from the opposite dimer via electrostatic interactions or hydrogen bonds at both ends of each respective β-sheet (Fig. 2C(i)-(ii)). Charge analysis in UCSF Chimera ^18^ shows that these two ends have high charge complementarity between dimers (Fig. 2C (iii)), suggesting possible hydrogen bonds or electrostatic interactions. Hydrophobic analysis shows that these β-sheets have an overall highly hydrophobic surface that also plays an important role in stabilizing their interactions.

We next compared the β-roll domain of iNS1 with the elongated β-sheet of the sNS1 stable tetramer. The iNS1 β-roll domain contains three β-strands (residues 1-7, 12-16 and 18-22), whereas we were unable to observe any density for residues 1-11 in the sNS1 stable tetramer, probably due to disorder in this region, and residues 15-24 form a long, continuous β-strand. This raises a question: is the sNS1 tetramer assembled from dimers of iNS1? If so, is it possible for the iNS1 β-roll structure to rearrange into the elongated β-sheet seen in the stable tetramer? If this were to happen, it would require the second β-strand of iNS1 to rotate by ∼120° with respect to the third β-strand to form the long, continuous β-strand we observed (Fig. 2D). Whether the iNS1 dimer is a precursor of the sNS1 stable tetramer is unknown. It is also possible that the sNS1 stable tetramer is assembled during translation of the NS1 protein by a pathway independent of iNS1 dimer formation, before being secreted.

### CryoEM structure of loose sNS1 tetramers

We also determined an 8Å resolution loose sNS1 tetramer structure (Figs. 3A(i) and Extended data Fig.4A) and by focused refinement on one of its constituent dimers, we obtained a 3.4Å resolution cryoEM map (Figs. 3A (ii) and Extended data Fig.4B (i-ii)). The overall structure of the dimer and its β-roll (Fig. 3A (ii)) is very similar to that of iNS1 (RMSD 0.8 Å), unlike the stable sNS1 tetramer. The overall 8Å resolution loose sNS1 tetramer structure shows that the two dimers are rotated by ∼35° with respect to each other and extra density is visible between the β-rolls of these two dimers (Fig. 3A(i)). Our high resolution sNS1 dimer structure within the loose tetramer also shows some strong extra densities above the β-roll (Fig. 3A(iii)). Previous work from other research groups ^13,14^ show that there are lipids associated with sNS1 that may help stabilize the overall structure, and which may contribute to these uninterpreted densities.

The difference between the stable and the loose tetramers is whether the residues near the N-terminus form a β-roll or an elongated β-sheet. Hence, this could be the determinant that defines stability.

An existing crystal structure of a ZIKV sNS1 ^11^ (PDB code: 5GS6), obtained by purifying sNS1 produced using a baculovirus expression system, is a dimeric structure. However, upon examination of the crystal packing in this structure, sNS1 appears to have a tetrameric structure similar to our loose tetramer (RMSD 4.2 Å) (Fig. 3A(iv)). This may suggest that at least one of the ZIKV sNS1 oligomers might be the loose tetramer structure.

### CryoEM structure of the sNS1 hexamer

We obtained an 8Å resolution sNS1 hexamer map (Figs. 3B(i) and Extended data Fig. 5A-B). The low resolution structure suggests that the NS1 molecules in the hexameric state are interacting loosely with each other. Fitting of iNS1 (Extended data Fig. 5C) suggests that three dimers interact with each other via their wing domain and the distal end of the β-ladder. This interaction is similar to that in the loose tetramer. The density in the central core of the hexamer is weak (Extended data Fig. 5B) and we were therefore unable to determine if the dimers have a β-roll or elongated β-sheet conformation. It is also possible that the interior of this hexamer contains lipid cargo, as suggested by other research groups ^13,14^ and this may prevent determination of the structure of this region.

The crystal structure of dengue dimeric iNS1 ^10^ shows that the crystal packing contains a trimer of dimers (i.e., hexamer) organization; however, there are very few interactions between the three dimers (Fig. 3B(ii)). Their positions in the crystal lattice are mainly supported by their interactions with other iNS1 molecules inside the crystal unit cell. This suggests that the hexameric packing of iNS1 may not be physiological. Comparison of our cryoEM sNS1 hexamer to this crystal hexamer packing shows a very different organization of dimers –they are rotated by ∼90° relative to each other (Fig. 3B(i) and (ii), right).

It is possible that the loose tetrameric structure is a transition intermediate of the hexameric structure (Fig. 3C). However, we observed loose tetramers forming ∼74% of the particle population (Fig. 1D), while hexamers only form ∼3%, suggesting that the loose tetramer may be a preferred or a lower energy oligomerization state.

It is possible that both the loose tetrameric and the hexameric structures are stabilized by lipids. To test this, we incubated the NS1 sample with detergent (0.05% DDM) and imaged it by cryoEM SPA. We observed that the percentage of loose tetramers and hexamers decreased dramatically after detergent treatment (Extended data Table 1), compared to untreated NS1 (Fig. 1E). This suggests that the lipids inside the loose tetramers and hexamers could be important for their structural integrity. A caveat to this conclusion is the detergent could also disrupt the hydrophobic interactions between the dimers.

### CryoEM structure of sNS1 complexed with Fab 5E3

We also investigated the cryoEM structure of sNS1 complexed with the Fab fragment of an anti-NS1 antibody 5E3. The 2D class averages (Fig. 4A) of the boxed particles from the cryoEM micrographs suggest that, when bound by Fabs, most sNS1 particles become dimeric. Unbound stable tetramers were also present (Fig. 4A) and formed ∼18% of the total particles. It suggests that the Fab 5E3 can readily dissociate the sNS1 loose tetrameric and hexameric structures. The stable tetrameric structure, on the other hand, is more resistant to Fab binding.

**Fig. 4.**
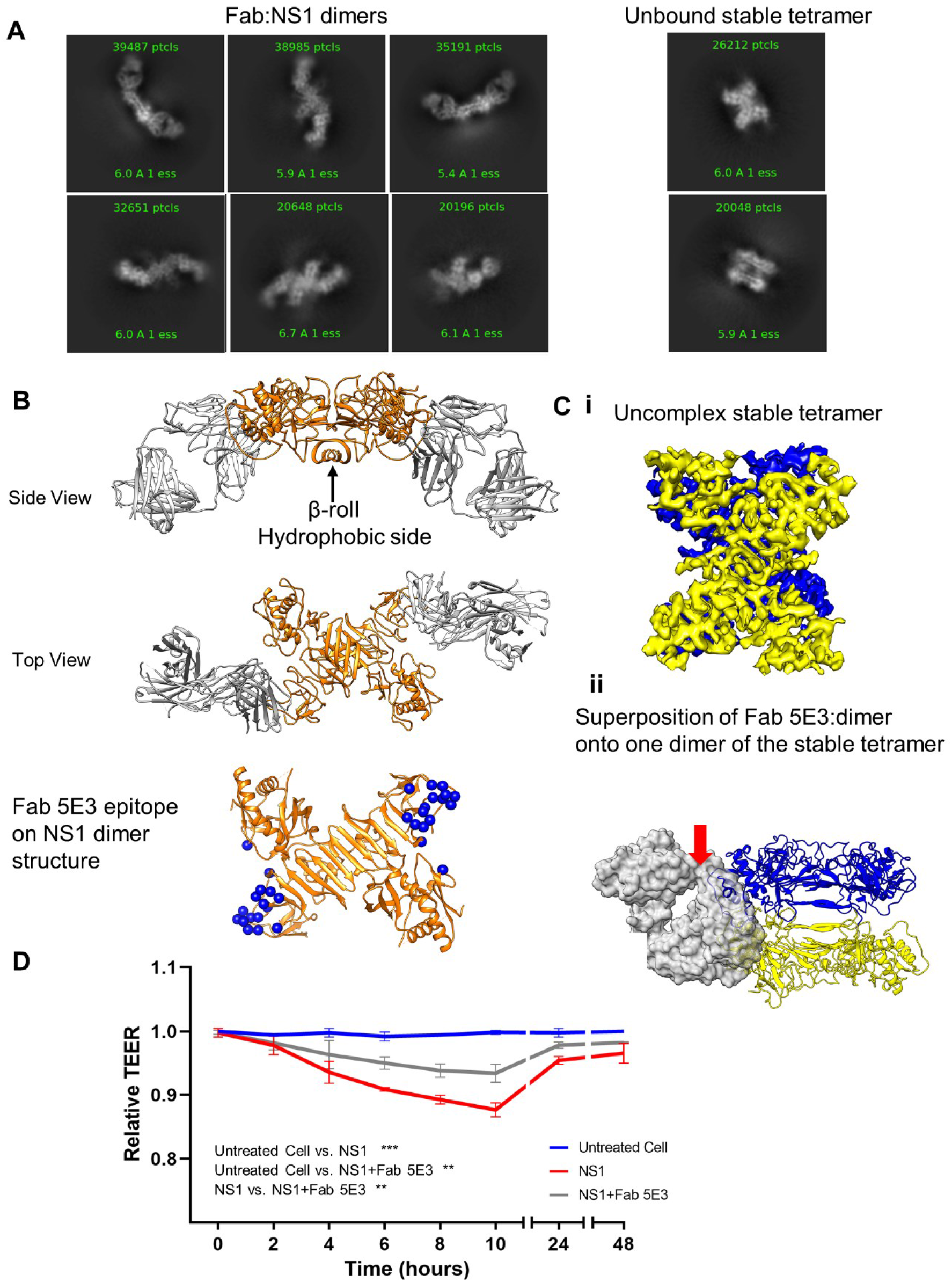
CryoEM structure of sNS1 complexed with Fab 5E3. **(A)** Some of the 2D class averages showing Fab bound dimers and also unbound stable tetramers. **(B)** The 3.5Å resolution structure of dimer NS1(orange) complexed with Fab 5E3 (grey) shows that they form an arch shape – the Fabs are bending towards the β-roll side of the sNS1 dimer. This NS1 dimer contains the β-roll structure which is also present in the loose tetramer and the hexamer. The Fabs bind to the ends of the β-ladder. The 5E3 epitope is shown as blue spheres. **(C) (i)** The 3.9Å resolution sNS1 unbound tetramer map. The two dimers of the tetramer are colored in blue and yellow. **(ii)** Superposition of Fab 5E3:dimer onto one dimer (yellow) of the stable tetramer shows that Fab (transparent grey surface) when bound will clash (red arrow) with the neighboring dimer (blue). **(D)** TEER assay showing Fab 5E3 can only partially prevent sNS1 from inducing human umbilical vein endothelial cell hyperpermeability. Data are the mean ± SEM from three independent experiments. Statistically significant differences between distinct groups (all time points combined for each group) compared to the untreated groups were determined by a two-way ANOVA analysis using Dunnett’s test for multiple comparisons. (***p < 0.001 and **p < 0.01).

The structure of the Fab 5E3:sNS1 dimer complex was determined to an overall resolution of 3.5Å (Fig. 4B, and Extended data Fig. 6A), as measured by gold standard FSC curve cutoff at 0.143 (Extended data Fig. 6B). The sNS1 dimer:Fab complex has an overall arch shape, where the Fabs bind to both ends of the β-ladders of the sNS1 dimer (Fig. 4B and Extended data Fig.6A) and are tilted towards the β-roll side of the sNS1.

Fab 5E3 interacts with the end of the β-ladder of NS1 (Fig. 4B, Extended data Figs. 6-7) via mainly the Fab heavy chain (Extended data Table 2). The 5E3 epitope is similar to two other previously published Fab:NS1 dimer complex ^19,20^ – Fabs 2B7 and 1G5.3 (Extended data Fig. 7). Comparison of these structures shows that these three Fabs bind at different angles to the NS1 β-ladder. It has been postulated previously^10^ that the hydrophobic β-roll may bind directly to cellular membranes (Extended data Fig. 1), which could be important for its pathogenesis, and therefore the binding orientation of Fab 5E3 may hinder this interaction. A similar neutralization mechanism was suggested for Fab 2B7 ^20^.

We observed that these NS1 dimers have β-roll structures (Figs. 4B and Extended data Fig.6C). This suggests that they are likely not derived from the stable tetramer which has an elongated β-sheet structure. Hence, they could be a broken down product of Fab binding to sNS1 loose tetramers (which contain β-roll structures) and perhaps also the hexamers. We superimposed the Fab:dimer complexes into the loose tetramer and hexamer, and they showed Fabs when bound, could clash with the neighbouring NS1 dimer (Extended data Fig. 8A-B). Analysis of the solvent accessibility of the epitopes on the loose tetramer (Extended data Fig. 8C) shows two of the epitopes (orange spheres) are partially exposed (37.5%), while the other two (cyan spheres), are fully exposed. All epitopes in the hexamer (Extended data Fig. 8D) are partially exposed (37.5%). Since Fab likely could bind to the loose tetramer and hexamer. This suggests that the loose tetramers and hexamers likely undergo motions that will at some time point, have their epitopes fully exposed for Fab binding, and once the Fab has bound, the loose tetramer and hexamers could be dismantled into dimers.

We reconstructed the unbound stable tetramer to 3.9Å resolution (Fig. 4C (i)), it is identical to the uncomplexed stable tetramer (Fig. 2). Analysis of the epitopes on all NS1 molecules in the unbound stable tetramer shows they are only partially exposed (37.5%) (Extended data Fig. 8E), thus discouraging Fab binding. Superposition of the Fab 5E3:dimer complex with one of the dimers in the stable tetramer structure (Fig. 4C (ii)) also shows that the Fab if bound, will clash with the wing domain of the neighbouring dimer. While the Fab is able to cause dissociation between dimers in the loose tetramer and hexamer, it is unable to do the same to the stable tetramer. This suggests that the two dimers in the stable tetramer have tighter interactions, thus disallowing antibody binding. The presence of unbound stable tetramer is consistent with the results of the TEER assay (Fig. 4D), showing Fab can only partially inhibit sNS1 induced endothelial cell hyperpermeability.

In conclusion, since the loose tetramer and hexamers are less stable, their quaternary structure can be easily disrupted by antibody binding. This may form part of the antibody neutralization mechanism. The stable tetramer is however more resistant to Fab 5E3 binding, and this may help sNS1 to evade the host immune response.

## Discussion

Previous studies ^13,14,21^ and our cross-linking experiments of sNS1 (Fig. 1B) consistently showed the presence of dimers, tetramers, and hexamers in semi- and fully denaturing SDS PAGE gel. Syzdykova *et al*. also showed by native gel that yellow fever virus sNS1 consists of mainly tetramers with little hexamers ^22^. Our cryoEM study shows the vast majority of DENV2 sNS1 particles exist as tetramers, while the hexamer only forms a minor population (Fig. 1E). Flamand and colleagues ^21^ performed small angle X-ray scattering (SAXS) experiments on sNS1 and observed particles with a homogenous gyration radius of 10nm (100 Å), which they concluded were likely hexamers. When we measured the dimensions of our cryoEM sNS1 tetrameric and hexameric structures (Extended data Fig. 2), their longest dimensions are the same (∼100 Å), thus correlating well with size of particles detected by the SAXS^21^. This suggests that SAXS may not be able to effectively differentiate between the heterogenous populations of the particles within the sNS1 sample.

NS1 plays multiple roles in dengue virus infection of mammalian cells and also mosquitoes. It was shown that sNS1 is one of the major factors contributing to the development of DHF/DSS in patients. Our cryoEM studies show structural details of this important sNS1 protein – (1) the quaternary organization of the different native oligomerization states, (2) how an antibody could bind and dismantle the loose tetrameric and hexameric sNS1 structures and also (3) how sNS1 stable tetramers could resist this antibody-induced disassembly. These findings will contribute to the design of therapeutics and vaccines against severe dengue disease.

## Methods

### sNS1 cloning, expression and purification

DENV 2 NS1-HisTag was cloned and expressed in Expi293 HEK cells. The sNS1 protein was purified by using a HisTrap column (GE Healthcare) and a gel-filtration chromatography. Protein purity was then assessed by SDS-PAGE.

### BS^3^ cross-linking of protein

BS^3^, a cross-linker, were added to NS1 in PBS to a final concentration of 5mM and the samples were incubated for 10, 20, 30 mins at room temperature. Reactions were stopped by adding Tris·HCl, pH 7.5 to a final concentration of 50mM and then the sample was added the SDS-PAGE loading buffer, boiled for 2 min and then analyzed on a 4–20 % SDS-PAGE gradient gels (Bio-Rad). The protein bands were then transferred to a nitrocellulose membrane, probed with anti-polyHistidine−peroxidase antibody and visualized by chemiluminescence.

### CryoEM sample preparation, image acquisition and reconstruction procedure

sNS1 was plunged frozen in liquid ethane by using the Vitrobot Mark IV plunger (FEI, Netherlands).

CryoEM micrographs of sNS1/ sNS1 complex were collected using a Titan Krios transmission electron microscope. Images were recorded by movie mode, and the frames from each movie were aligned using MotionCor2 ^23^ to produce full dose images. The astigmatic CTF parameters were estimated and accounted for during orientation search. A total of 2,342,754 particles were picked. 2D class averages of particles were done. We then performed 3D classification and reconstruction using C1 symmetry using the program Relion ^15^ to remove broken and distorted particles. For the class with stable tetramers, we performed classification and refinement by imposing D2 symmetry. For the loose tetramers, we reconstructed the whole particle using C1 symmetry and also performed local refinement on one of its dimers. For the hexamers, we imposed D3 symmetry. Resolution was determined by using gold standard Fourier shell correlation and local map resolution was estimated with ResMap ^25^.

### Protein structure building

The sNS1 structures were first interpreted by approximate fitting of the NS1 dimeric crystal structure using “fit-in-map” function in Chimera ^18^. We then fit individual residues into densities ^26^. We then refined the structures using the “phenix.real_space_refine” procedure in the Phenix software package, with default parameters and rigid body refinement using secondary-structure and torsion angle restraints ^27^. The final coordinates containing an asymmetric unit (i.e., a protomer in the stable tetramer structure) were then used to build the biological unit (full tetramer) by using the command “sym” in UCSF Chimera ^18^. The final coordinates of the asymmetric units were checked using MolProbity ^28^. Maps and structures shown in the Figs were generated by using UCSF Chimera and Coot.

### TEER assay

A monolayer of Human Umbilical Vein endothelial cells (80%–90% confluency) cultured in T75 flasks were detached using trypsin-EDTA (0.25%). The cells were then resuspended using fresh culture media and then counted using an automated cell counter. 60,000 cells (300 μL per well) were seeded onto the apical side of Transwell inserts. Each Transwell was transferred inside a 24-well format plate containing 1.5 mL of endothelial cell culture media. Transwells containing endothelial cells were incubated at 37°C and 5% CO2 for 3∼4 days, and 50% of the culture medium was changed in each well 48 hours post-seeding. Cells were grown until the TEER values reached ∼ 160 to 170 Ohms (Ω), indicating 100% cell confluency. After this, 3 μg NS1 protein / NS1_Fab5E3 complex (molar ratio 1:1.2) was added to the apical side of the Transwell insert. Electrical resistance values, measured in Ohms (Ω), were recorded at every 2-hour time-points following the addition of the proteins using an Epithelial Volt Ohm Meter with the “chopstick” electrodes. Endothelial permeability was expressed as relative TEER which represents a ratio of resistance values (Ω) as follows: (Ω experimental condition - Ω medium alone)/(Ω nontreated endothelial cells - Ω medium alone).

## Acknowledgments

We thank Valerie S-Y Chew, Xinni Lim and Qunfei Zhou for help in experiments. This work was supported by Duke-NUS Signature Research Programme funded by the Ministry of Health, Singapore and National Research Foundation Investigatorship award (NRF-NRFI2016-01) awarded to SML. AWKT is supported by a Khoo Postdoctoral Fellowship Award (Duke-NUS-KPFA/2021/0044) from Duke-NUS Medical School and the Estate of Tan Sri Khoo Teck Puat.

## Author information

### Contributions

S.M.L. supervised the project. S.M.L. and B.S. designed research studies; J.S.G.O. produced sNS1. W.D., J.M. and G.S. produced Fab. B.S. and J.S.G.O. conducted experiments. T.S.N., A.W.K.T. and J.S. collected cryo-EM data; S.M.L., B.S., G.F. and V.A.K. analyzed data; S.M.L. and B.S. wrote the manuscript.

### Corresponding author

Correspondence to Shee-Mei Lok.

## Ethics declarations

### Competing interests

The authors declare no competing interests.

## Additional information

**Extended Data Fig.1.**
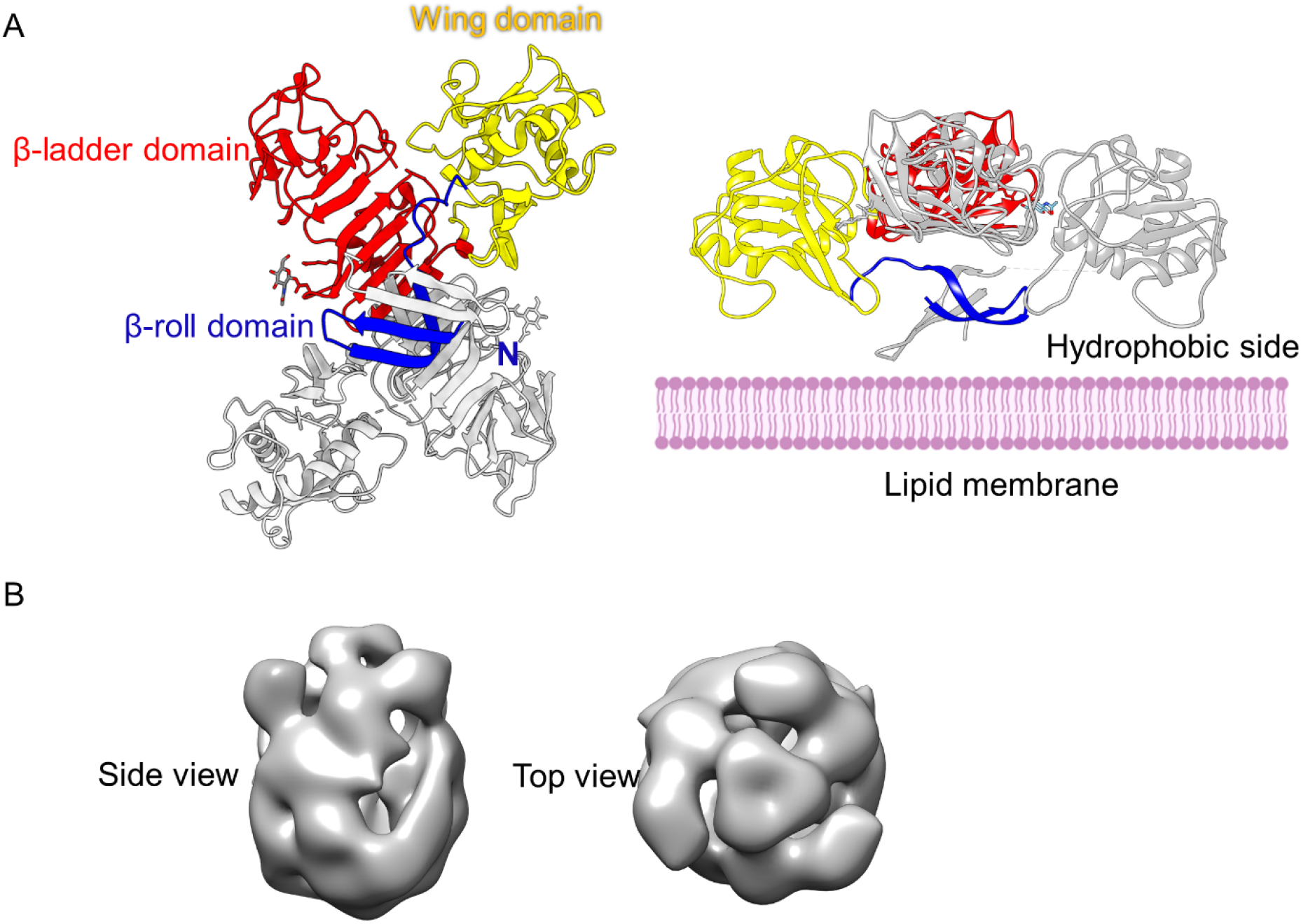
Previously published structures of the dengue NS1. (A) Crystal structure of dengue dimeric iNS1 (PDB:5GS6). Left: top view, looking from the β-roll of the iNS1 dimer. One protomer contains three domains: starting from the N-terminus-β-roll (blue), wing domain (yellow) and the β-ladder (red). It interacts with the other protomer (colored in grey) through extensive hydrogen bonds. Right: side view of the iNS1 dimer. The β-roll is highly hydrophobic and is predicted to interact with the cellular membrane. (B) A previously published low resolution EM reconstruction of the sNS1 (EMDB:2073). This map is reconstructed from negatively stained sNS1 and C3 symmetry was imposed ^13^.

**Extended Data Fig.2.**
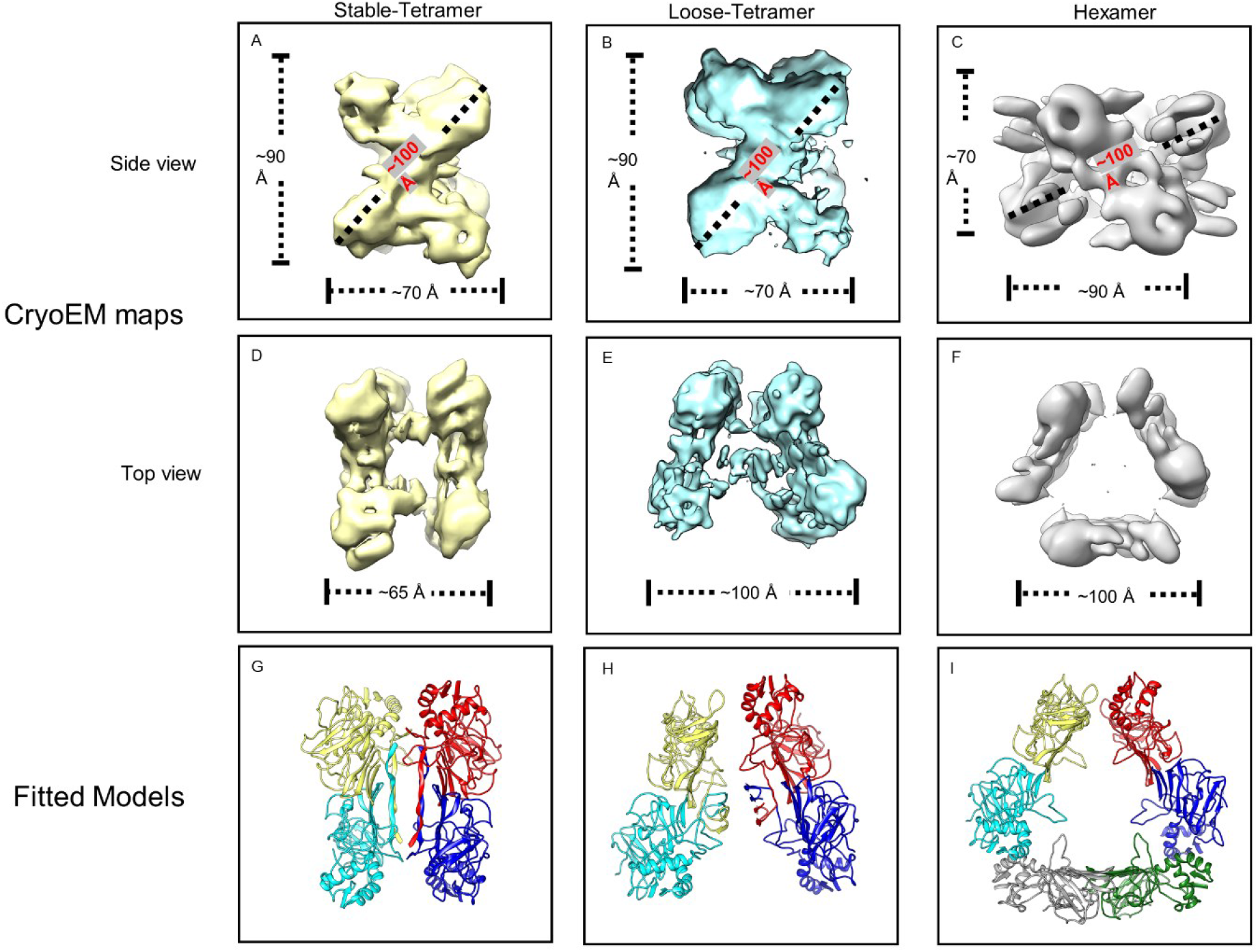
The initial sNS1 cryoEM maps after 3D classification using C1 symmetry. Dimensions of the stable, loose tetramers and hexamers are shown. Fitted models are shown on the bottom row.

**Extended Data Fig.3.**
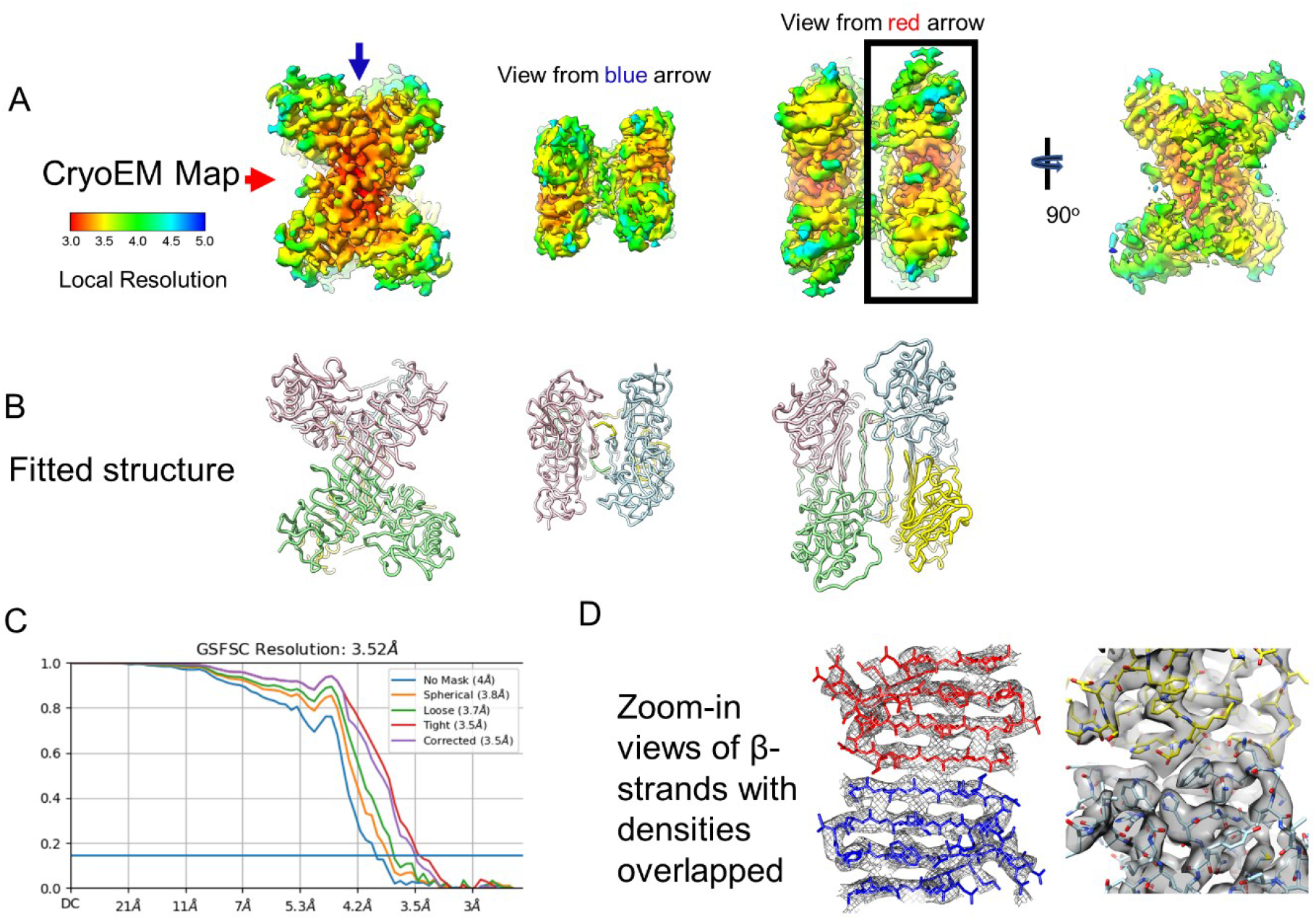
The 3.5Å resolution sNS1 stable tetrameric structure. (A) Different views of the cryoEM map displayed at high contour level. The densities are colored according to their local resolution. (B) The fitted structure show in the same view as the cryoEM map directly above (in (A)). There are two dimers facing each other. One dimer is colored in pink and green, while the other is light blue and yellow. (C) Fourier shell correlation plot showing resolution. (D) Zoom-in views of density map showing the β-strands densities are well resolved (left). Some of the side chains densities (right) can also be observed.

**Extended Data Fig.4.**
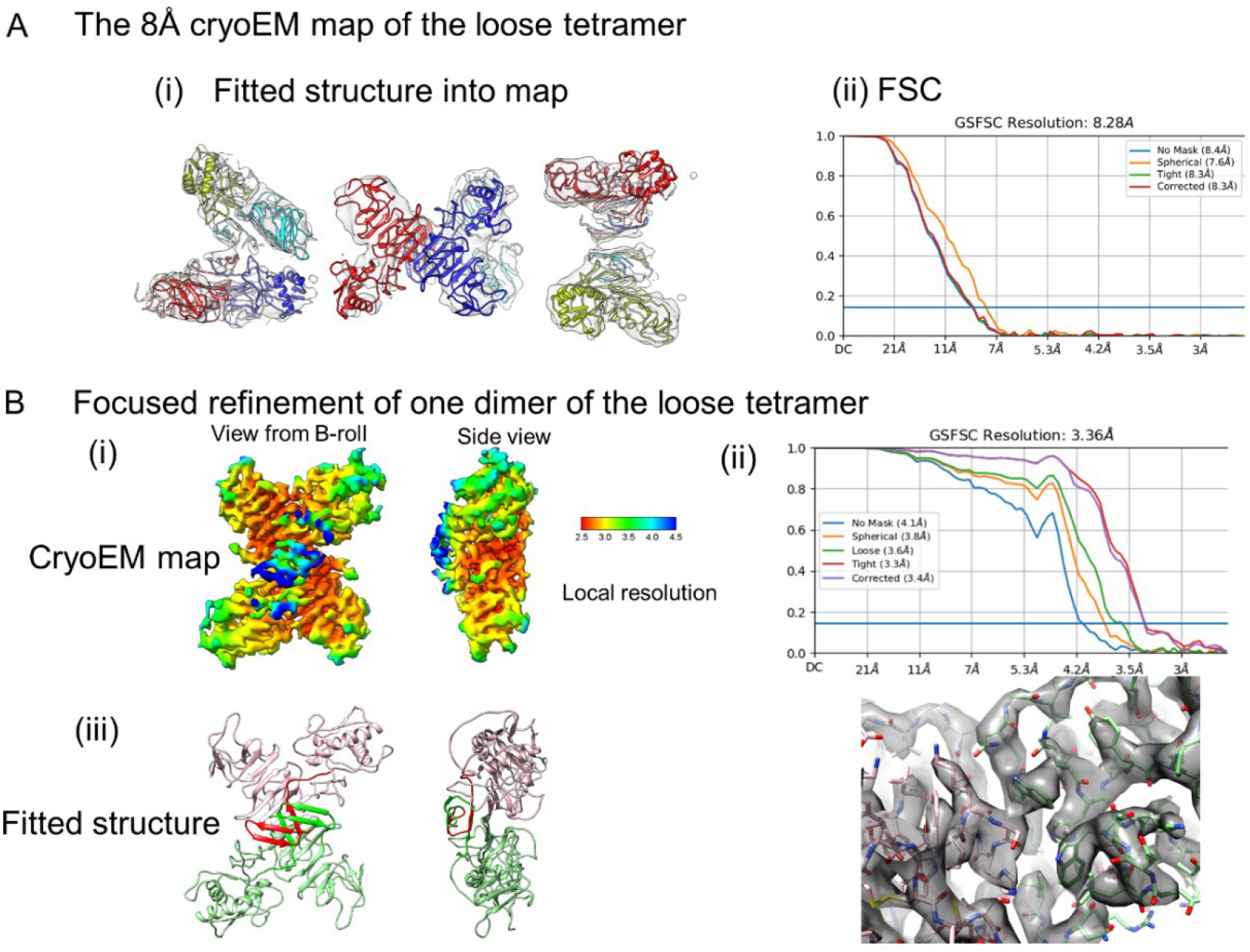
CryoEM maps of the sNS1 loose tetramer. (A) 8Å cryoEM map of the loose tetramer. (i) Different views of the fitted structure of the loose tetramer into the density map (transparent grey). (ii) FSC plot showing resolution. (B) The 3.4Å cryoEM map of one dimer of the loose tetramer after focused refinement. (i) Different views of the cryoEM map. Density is colored according to local resolution. (ii) Up: FSC plot showing resolution of the map: Down: zoom-in view showing that some side chain densities are well resolved. (iii) The fitted structure of one of the dimers (pink and light green) is very similar to that of dengue iNS1. The β-roll in each protomer is colored in a darker shade of the respective protomer’s color.

**Extended Data Fig.5.**
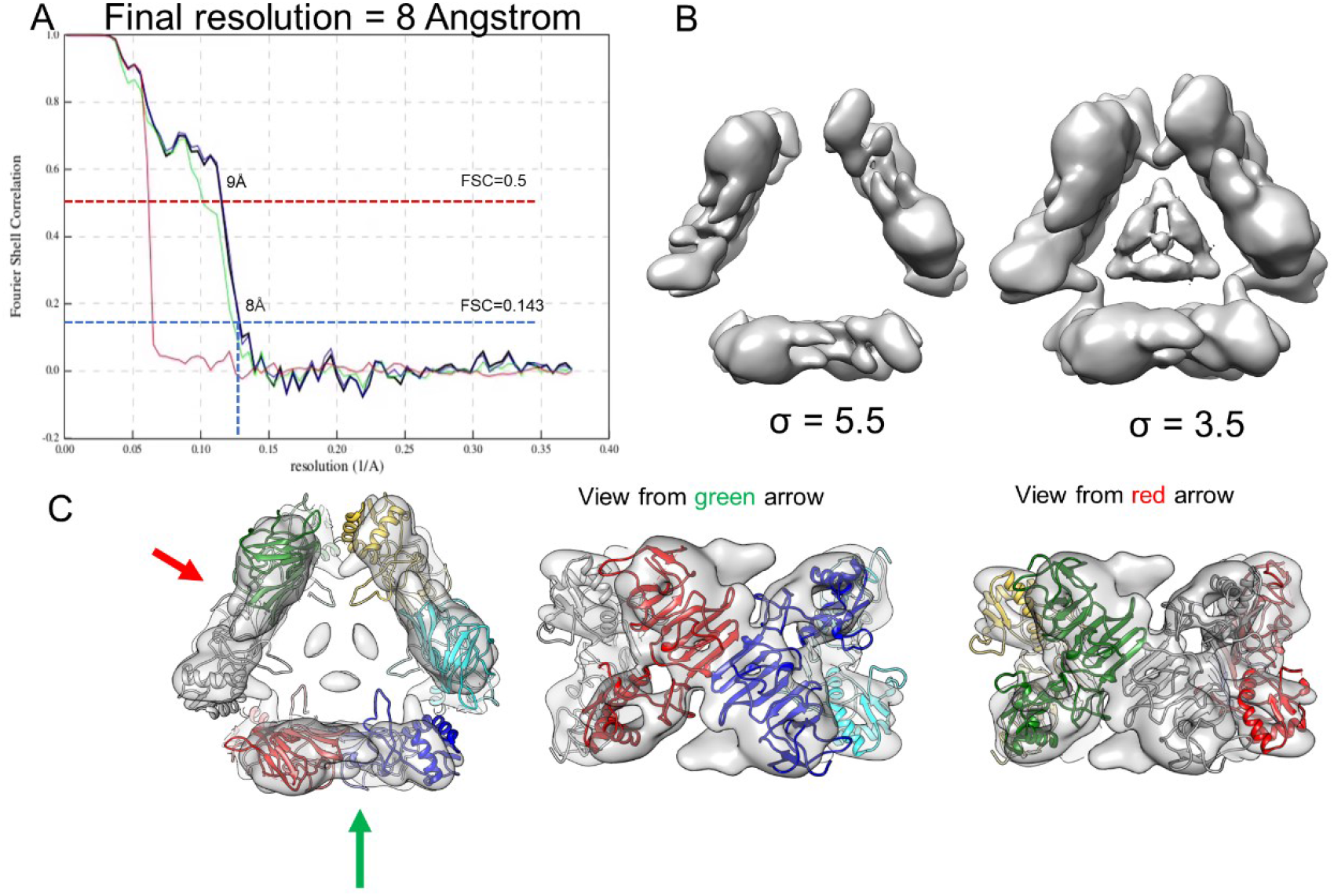
The 8Å cryoEM map of the sNS1 hexamer. (A) FSC plot showing resolution of the map. (B) The hexamer density displayed at different contour levels. At lower contour levels, densities at the interior of the hexamer can be seen. (C) Different views of the fit of the three dimers into the hexamer density map.

**Extended Data Fig.6.**
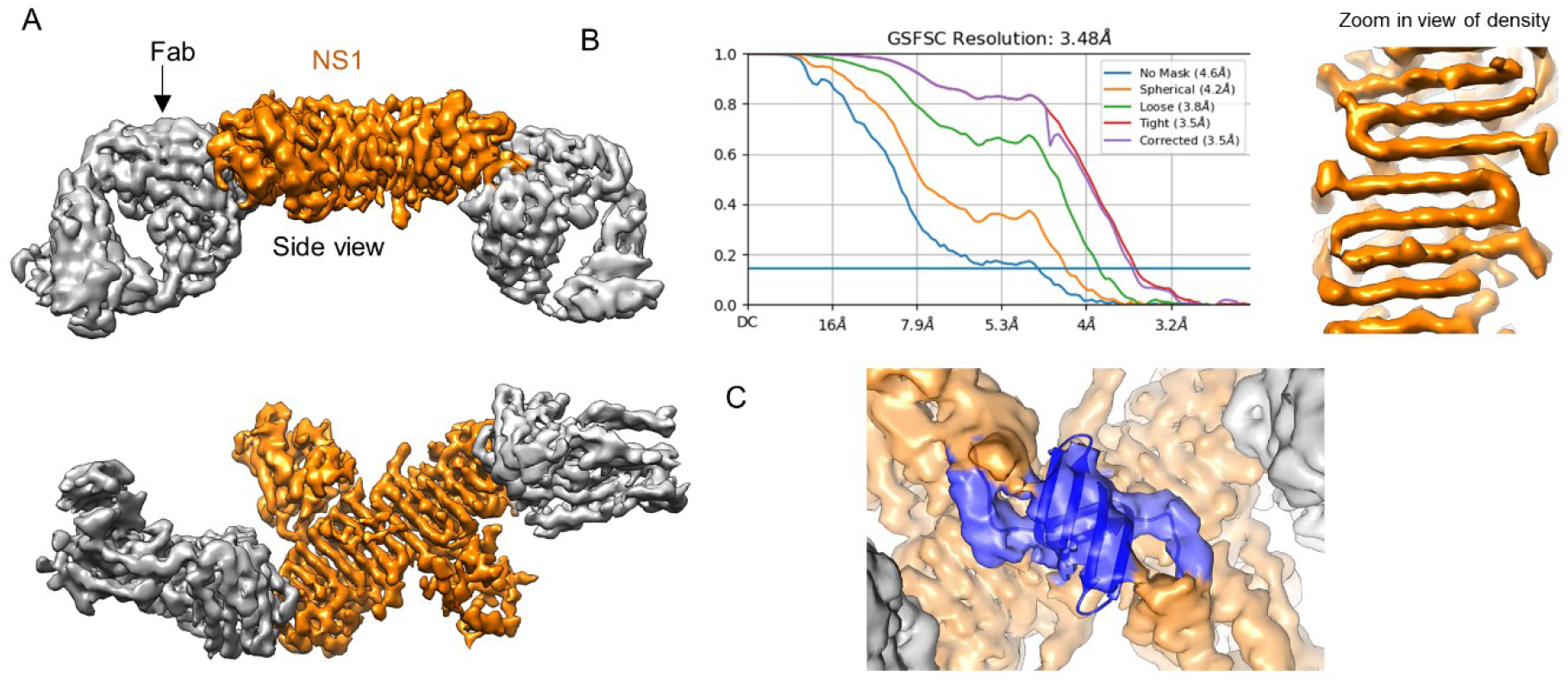
3.5Å resolution cryoEM structure of Fab 5E3:sNS1 dimer. (A) Side and top views of the cryoEM map of the Fab 5E3:sNS1 dimer. NS1 is colored in orange while Fabs in grey. (B) Left: FSC plot showing the resolution of the map. Right: zoom-in view showing well-resolved β-strands. (C) Zoom-in view of the fitted β-roll into its corresponding density.

**Extended Data Fig.7.**
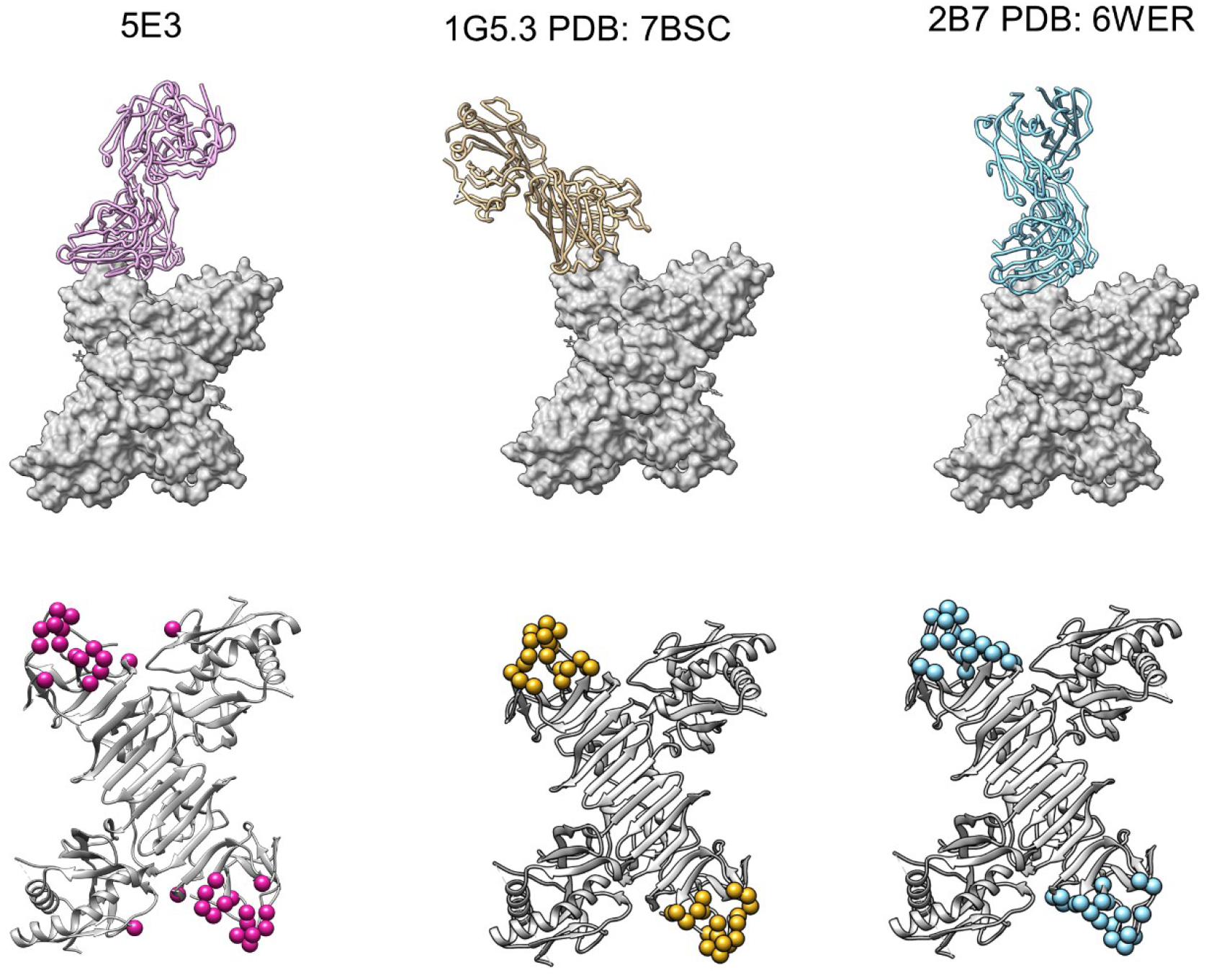
Comparison of the binding (top panels) and epitope (bottom panels) of Fab 5E3 to other previously published Fab:NS1 complexes ^19,20^.

**Extended Data Fig.8.**
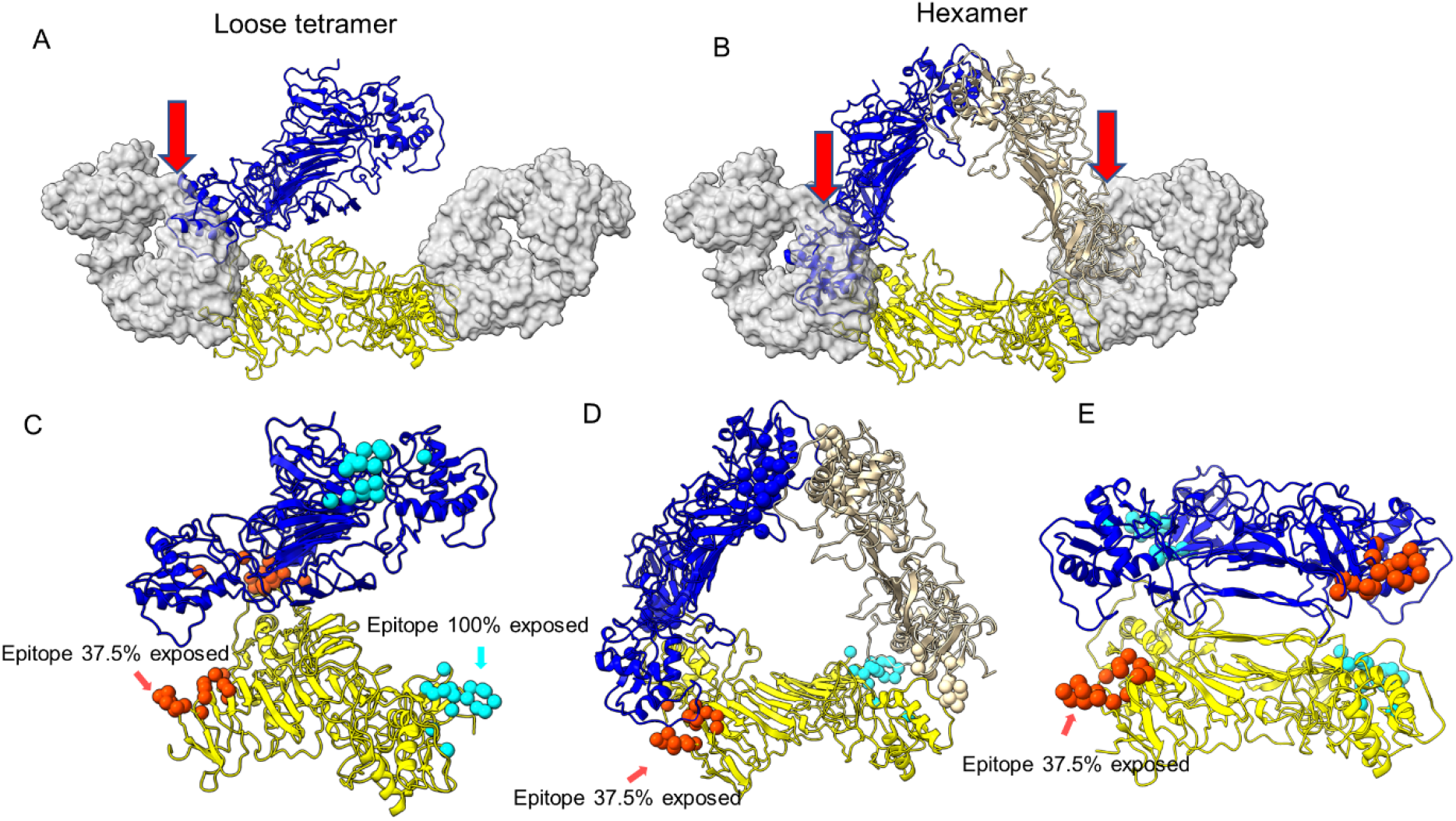
Superposition of Fab 5E3:dimer into the (A) loose tetramer and (B) hexamer and the analysis of percentage exposure of epitopes (C-E) in the stable, loose tetramers and hexamers. (A-B) Superposition of Fab 5E3:dimer onto one dimer (yellow) of the loose tetramer and hexamer. Fab is shown as transparent grey surface and clashes with neighboribg NS1 dimer is indicated with red arrows. (C-E) The percentage exposure of epitopes in the oligomers. The two epitopes within one dimer are colored in orange and cyan. (C) In the loose tetramer, 37.5% of the orange epitope is exposed while the cyan epitope is fully exposed (100%). (D) In the hexamer, 37.5% of both the orange and cyan epitopes are exposed. (E) In the stable tetramer, the two epitopes are both 37.5% exposed.

**Extended Data Table 1.** The percentage of particles in different oligomerization states determined after 3D classification of NS1 treated with detergent (0.05% DDM).

|  | Stable Tetramer | Loose Tetramer | Hexamer |
| --- | --- | --- | --- |
| Particles | 126130 | 90928 | 0 |
| Percentage | 58% | 42% | 0 % |

**Extended Data Table S2.** List of hydrogen bonds/electrostatic interactions between Fab 5E3 (heavy and light chains) with the NS1 protein. The interacting CDRs of the Fabs are indicated.

| **NS1** | **Fab Heavy Chain** | **Fab Light Chain** |
| --- | --- | --- |
| **138** |  | **27(CDR1)** |
| **269** |  | **94(CDR3)** |
| **281** | **52/53/54/55/56(CDR2)** |  |
| **299** | **102/103(CDR3)** |  |
| **301** | **33(CDR1)** |  |
| **302** | **103(CDR3)** |  |
| **303** | **103(CDR3)** |  |
| **304** | **32/106(CDR1/CDR3)** |  |
| **305** | **31/33/103(CDR1/CDR3)** |  |
| **306** | **31(CDR1)** |  |
| **307** | **31/53(CDR1/CDR2)** |  |
| **324** | **56(CDR2)** |  |
| **325** | **57(CDR2)** |  |
| **326** | **57/102/100(CDR2/CDR3)** |  |
| **327** | **100/102(CDR3)** | **94(CDR3)** |
| **328** | **100(CDR3)** |  |

## Notes

### Competing Interest Statement

The authors have declared no competing interest.

## References

1 Gubler, D. J. Dengue and dengue hemorrhagic fever. Clin Microbiol Rev 11, 480–496, doi:10.1128/CMR.11.3.480 (1998).

2 Thomas, S. J. & Yoon, I. K. A review of Dengvaxia(R): development to deployment. Hum Vaccin Immunother 15, 2295–2314, doi:10.1080/21645515.2019.1658503 (2019).

3 Halstead, S. B. Neutralization and antibody-dependent enhancement of dengue viruses. Adv Virus Res 60, 421–467, doi:10.1016/s0065-3527(03)60011-4 (2003).

4 Beatty, P. R. et al. Dengue virus NS1 triggers endothelial permeability and vascular leak that is prevented by NS1 vaccination. Sci Transl Med 7, 304ra141, doi:10.1126/scitranslmed.aaa3787 (2015).

5 Carpio, K. L. & Barrett, A. D. T. Flavivirus NS1 and Its Potential in Vaccine Development. Vaccines (Basel) 9, doi:10.3390/vaccines9060622 (2021).

6 Modhiran, N. et al. Dengue virus NS1 protein activates immune cells via TLR4 but not TLR2 or TLR6. Immunol Cell Biol 95, 491–495, doi:10.1038/icb.2017.5 (2017).

7 Puerta-Guardo, H., Glasner, D. R. & Harris, E. Dengue Virus NS1 Disrupts the Endothelial Glycocalyx, Leading to Hyperpermeability. PLoS Pathog 12, e1005738, doi:10.1371/journal.ppat.1005738 (2016).

8 Libraty, D. H. et al. High circulating levels of the dengue virus nonstructural protein NS1 early in dengue illness correlate with the development of dengue hemorrhagic fever. J Infect Dis 186, 1165–1168, doi:10.1086/343813 (2002).

9 Liu, J. et al. Flavivirus NS1 protein in infected host sera enhances viral acquisition by mosquitoes. Nat Microbiol 1, 16087, doi:10.1038/nmicrobiol.2016.87 (2016).

10 Akey, D. L. et al. Flavivirus NS1 structures reveal surfaces for associations with membranes and the immune system. Science 343, 881–885, doi:10.1126/science.1247749 (2014).

11 Xu, X. et al. Contribution of intertwined loop to membrane association revealed by Zika virus full-length NS1 structure. EMBO J 35, 2170–2178, doi:10.15252/embj.201695290 (2016).

12 Wang, D. et al. A Mutation Identified in Neonatal Microcephaly Destabilizes Zika Virus NS1 Assembly in Vitro. Sci Rep 7, 42580, doi:10.1038/srep42580 (2017).

13 Muller, D. A. et al. Structure of the dengue virus glycoprotein non-structural protein 1 by electron microscopy and single-particle analysis. J Gen Virol 93, 771–779, doi:10.1099/vir.0.039321-0 (2012).

14 Gutsche, I. et al. Secreted dengue virus nonstructural protein NS1 is an atypical barrel-shaped high-density lipoprotein. Proc Natl Acad Sci U S A 108, 8003–8008, doi:10.1073/pnas.1017338108 (2011).

15 Scheres, S. H. RELION: implementation of a Bayesian approach to cryo-EM structure determination. J Struct Biol 180, 519–530, doi:10.1016/j.jsb.2012.09.006 (2012).

16 Punjani, A., Rubinstein, J. L., Fleet, D. J. & Brubaker, M. A. cryoSPARC: algorithms for rapid unsupervised cryo-EM structure determination. Nat Methods 14, 290–296, doi:10.1038/nmeth.4169 (2017).

17 Punjani, A., Zhang, H. & Fleet, D. J. Non-uniform refinement: adaptive regularization improves single-particle cryo-EM reconstruction. Nat Methods 17, 1214–1221, doi:10.1038/s41592-020-00990-8 (2020).

18 Pettersen, E. F. et al. UCSF Chimera--a visualization system for exploratory research and analysis. J Comput Chem 25, 1605–1612, doi:10.1002/jcc.20084 (2004).

19 Modhiran, N. et al. A broadly protective antibody that targets the flavivirus NS1 protein. Science 371, 190–194, doi:10.1126/science.abb9425 (2021).

20 Biering, S. B. et al. Structural basis for antibody inhibition of flavivirus NS1-triggered endothelial dysfunction. Science 371, 194–200, doi:10.1126/science.abc0476 (2021).

21 Flamand, M. et al. Dengue virus type 1 nonstructural glycoprotein NS1 is secreted from mammalian cells as a soluble hexamer in a glycosylation-dependent fashion. J Virol 73, 6104–6110, doi:10.1128/JVI.73.7.6104-6110.1999 (1999).

22 Syzdykova, L. R. et al. Fluorescent tagging the NS1 protein in yellow fever virus: Replication-capable viruses which produce the secretory GFP-NS1 fusion protein. Virus Res 294, 198291, doi:10.1016/j.virusres.2020.198291 (2021).

23 Zheng, S. Q. et al. MotionCor2: anisotropic correction of beam-induced motion for improved cryo-electron microscopy. Nat Methods 14, 331–332, doi:10.1038/nmeth.4193 (2017).

24 Tegunov, D. & Cramer, P. Real-time cryo-electron microscopy data preprocessing with Warp. Nat Methods 16, 1146–1152, doi:10.1038/s41592-019-0580-y (2019).

25 Kucukelbir, A., Sigworth, F. J. & Tagare, H. D. Quantifying the local resolution of cryo-EM density maps. Nat Methods 11, 63–65, doi:10.1038/nmeth.2727 (2014).

26 Emsley, P. & Cowtan, K. Coot: model-building tools for molecular graphics. Acta Crystallogr D Biol Crystallogr 60, 2126–2132, doi:10.1107/S0907444904019158 (2004).

27 Adams, P. D. et al. PHENIX: a comprehensive Python-based system for macromolecular structure solution. Acta Crystallogr D Biol Crystallogr 66, 213–221, doi:10.1107/S0907444909052925 (2010).

28 Williams, C. J. et al. MolProbity: More and better reference data for improved all-atom structure validation. Protein Sci 27, 293–315, doi:10.1002/pro.3330 (2018).

